# epsSMASH uncovers exopolysaccharide biosynthetic gene clusters in environmental and human microbiomes

**DOI:** 10.64898/2025.12.21.693542

**Authors:** Anders Ogechi Hostrup Daugberg, Angie Waldisperg, Marie Riisgaard-Jensen, Sofie Zacho Vestergaard, Roberto Sánchez Navarro, Tilmann Weber, Kai Blin, Simon Shaw, Per Halkjær Nielsen, Morten Kam Dahl Dueholm

**Author notes:** **Correspondence:** Morten Kam Dahl Dueholm, Center for Microbial Communities, Department of Chemistry and Bioscience, Aalborg University, Fredrik Bajers Vej 7H, 9220 Aalborg, Denmark.

## Abstract

Biofilms represent the default mode of bacterial life in natural and built environments, with extracellular polysaccharides (exoPS) serving as essential structural and functional components of the biofilm matrix. Despite their importance, exoPS production in these environments is largely unknown. Here we present epsSMASH, a bioinformatic tool and web service for predicting known and novel exoPS biosynthetic gene clusters (BGCs) in bacterial genomes. Benchmarking showed that comprehensive detection of exoPS gene clusters requires highly contiguous high-quality genome assemblies. We applied epsSMASH to high-quality bacterial genome catalogues representing four major ecosystems: Human gut, soil, ocean and activated sludge from wastewater treatment systems. In all catalogues, epsSMASH identified exoPS BGCs in most genomes (52.8-85.4%), with a median of 1-2 exoPS BGCs per genome. The number of exoPS BGC per genome was highly variable, with some taxa containing up to 19 distinct exoPS BGCs. Pel BGCs were abundant in human gut, ocean and activated sludge microbiomes, and were detected in 14 different phyla, making it the most phylogenetically widespread BGC in these environments. The vast majority (62-96%) of detected exoPS BGCs were uncharacterised. By constructing gene cluster families from uncharacterised systems, we identified novel and phylogenetically widespread exoPS BGCs. We investigated a novel exoPS gene cluster from the activated sludge microbiome and showed that it is conserved in most genera within the order Sphingomonadales. Our results highlight the remarkable number of uncharacterised exoPS gene clusters in environmental microbiomes and establish epsSMASH as an effective tool for identifying and classifying novel exoPS systems.

## Introduction

Secretion of polysaccharides into the extracellular space is a fundamental and widespread microbial trait that underpins key biological functions such as biofilm formation, host interaction, and protection against environmental stress^1^. Lipopolysaccharides (LPS) are constituents of the extracellular layer of the Gram-negative outer membrane and enable the evasion of host cell defences^2^. Capsular polysaccharides (CPS) envelop both Gram-positive and Gram-negative bacteria in capsular structures, protecting them from toxins and host immune responses as well as assisting in attachment and aggregation^3^. Some secreted polysaccharides are less strongly associated with the cell membrane and end up in the extracellular space. These play a key role in all stages of the biofilm lifecycle from initial attachment until dispersal of the mature biofilm, providing antimicrobial resistance, viscoelasticity and structural integrity^4^. In this study we use the term “exopolysaccharides” (exoPS) to refer to all polysaccharides that are secreted into the extracellular space, regardless of their attachment to the polysaccharide-producing cell.

exoPS biosynthesis can be grouped into four major pathways: i) the synthase-dependent, ii) the sucrase-dependent, iii) the ABC transporter-dependent, and iv) the Wzx/Wzy-dependent pathways^1,5^. All known synthase- and sucrase-dependent pathways produce exoPS which are not covalently attached to the cell surface^1^. The ABC transporter-dependent pathways produce either CPS or LPS, while the Wzx/Wzy-dependent pathways are capable of producing CPS, LPS, as well as exoPS which are not associated with the cell surface, such as Psl and xanthan^6^. With the exception of the sucrase-dependent pathways, which are dependent on a single extracellular sucrase, the genes required for exoPS biosynthesis are colocalised in biosynthetic gene clusters (BGCs)^6^.

Knowledge about the colocalisation of genes involved in the same biosynthetic pathway has been used in the past to predict gene clusters responsible for bacterial secondary metabolite production^7^, primary metabolite production in the human gut microbiome^8^, and bacterial catabolism in the rhizosphere^9^. Requiring colocalisation of genes in addition to sequence similarity provides an additional safeguard against false-positive BGC predictions^10^. A well-known example of this approach is antiSMASH, which uses profile Hidden Markov Models (pHMMs) to detect individual genes and manually curated BGC rules to predict the genetic potential for specialised/secondary metabolites^7^.

Despite the ubiquity of exoPS, only a few studies have investigated the genomic potential of exoPS production in environmental and human microbiomes^11,12^. Here we present epsSMASH, a bioinformatic tool based on the antiSMASH framework that can detect known and previously undescribed exoPS BGCs in bacterial genomes. To assess the genomic potential for exoPS production in different ecosystems, we apply epsSMASH to four high-quality genome databases representing the microbiome of the human gut, soil, ocean and activated sludge from wastewater treatment systems.

## Results

epsSMASH is a python-based bioinformatic tool that detects exoPS BGCs in bacterial genome sequences. It is a customised version of antiSMASH 7.0^13^ where secondary metabolite pHMMs and detection rules have been substituted for ones specific to exoPS BGCs. epsSMASH accepts amino acid sequence files in the GBK, EMBL and FASTA formats as input (**Figure 1a**). If a nucleotide FASTA file is used as input, epsSMASH will first predict genes using Prodigal (v2.6.3)^14^. Using HMMER v3.4^15^, each gene is searched against a collection of pHMMs which are each specifically built to detect a certain type of exoPS BGC (**Supplementary Note 1**).

**Figure 1:**
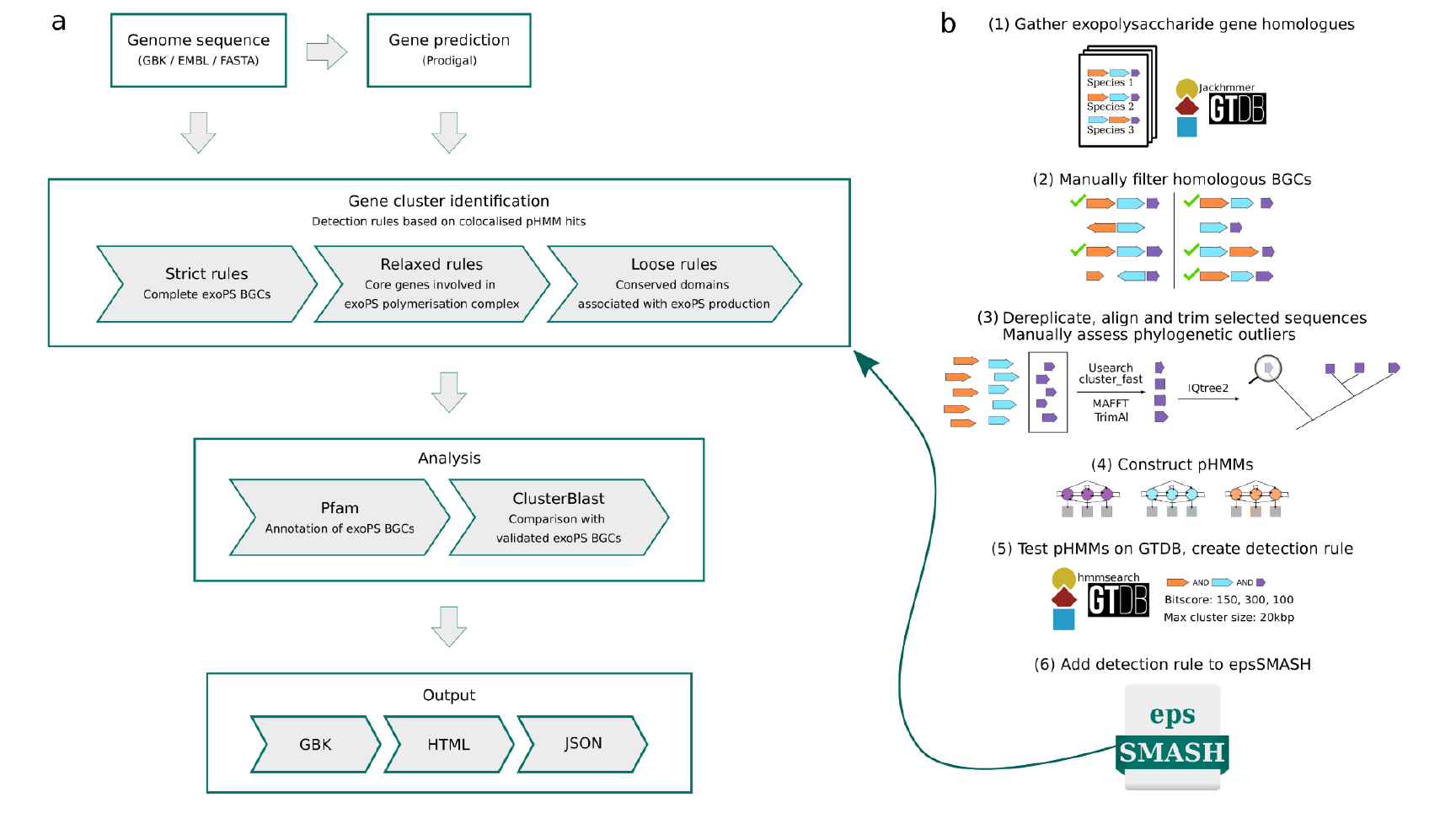
epsSMASH and epsProtocol workflows. **a**. The epsSMASH exoPS BGC genome mining workflow. **b**. epsProtocol workflow. (1) A set of known and characterised exoPS BGCs were curated from literature and each gene was searched against the Genome Taxonomy Database using Jackhmmer. (2) For each characterised exoPS BGC, the hits from the Jackhmmer search were grouped based on a minimum intergene distance of 5000 bp. These putative gene clusters were manually filtered to match the synteny and gene composition described in literature, resulting in a set of validated exoPS BGCs. (3) Each set of genes in the set of validated exoPS BGC are dereplicated, aligned and trimmed. Phylogenetic trees are made for each gene set and phylogenetic outliers are removed based on manual inspection. (4) profile Hidden Markov Models (pHMMs) are created for each set of genes. (5) The newly created pHMMs are searched against the Genome Taxonomy Database using hmmsearch, and the hits are grouped based on a minimum intergene distance criteria of 5000 bp. One or more epsSMASH rules are created using antiSMASH rule syntax. Bit score thresholds and maximum cluster size are set for each pHMM to improve rule accuracy (Numbers in the figure are examples). (6) The newly created rule and the related pHMMs are added to epsSMASH.

Subsequently, a set of manually curated epsSMASH rules identify exoPS BGCs in the sequence based on the colocalisation of specific pHMM hits. The most conservative of these rules require that the detected exoPS BGCs share the gene composition of the representative exoPS BGC in literature. We refer to these as “strict” rules.

To allow for the detection of alternative forms of known exoPS BGC or previously undescribed exoPS BGCs, we also created sets of “relaxed” and “loose” rules (**Supplementary Note 1**).

Relaxed rules are less stringent versions of strict exoPS synthase-dependent rules, which only require core biosynthetic genes to be present. The loose rules were created to search for uncharacterised Wzx/Wzy- and ABC transporter-dependent exoPS BGCs. These require only the genetic components for biosynthesis of activated sugars and the exoPS secretion system to be present. By default, epsSMASH uses strict, relaxed and loose rules, but one or more rulesets can be disabled in the command-line version. To assist BGC analysis, epsSMASH includes an option to annotate detected gene clusters with Pfam domains. Additionally, “ClusterBlast” can be enabled, which compares detected exoPS BGCs with a database of manually validated gene clusters. All epsSMASH results are saved in a user-friendly HTML page, which can be used to analyse the detected gene clusters, as well as in JSON and GBK files for downstream analysis.

To develop the rulesets for identifying exoPS BGCs, we manually screened the literature and gathered a set of 28 known exoPS BGCs, which had at least one peer-reviewed article describing their gene compositions and synteny with accompanying NCBI accession numbers (**Supplementary Table 1**). These included 14 synthase-dependent, 2 sucrase-dependent, 1 ABC transporter-dependent, and 11 Wzx/Wzy-dependent systems. To ensure that epsSMASH rules were created in a rigorous and reproducible manner, we developed a standardised, Snakemake-based workflow, epsProtocol (**Figure 1b**, see **Supplementary Note 1** for details). epsProtocol finds sequence homologs for each translated gene of a specific exoPS BGC from literature in the Genome Taxonomy Database (GTDB, v220)^16^. The detected homologs were clustered into putative exoPS BGCs if their intergene distances are below 5000 bp. These exoPS BGCs were manually validated to filter out any exoPS BGCs which do not contain core biosynthetic genes or have a different synteny from what is described in the literature. The translated gene sequences from these manually validated exoPS BGCs were clustered based on 90% average amino acid identity, aligned and trimmed before phylogenetic trees for each translated gene were created to filter out any obvious outliers. Finally, pHMMs for each gene in the exoPS BGC were constructed from these alignments.

**Table 1:** Statistics and epsSMASH results for the four HQ MAG databases.

| Genome catalogue | HQ MAGs | Median contig N50 (IQR) | exoPS BGCs | MAGs with exoPS BGCs (%) | Median BGCs per MAGs (IQR) |
| --- | --- | --- | --- | --- | --- |
| Microflora Danica | 4,894 | 388.4 kbp (226.3 - 691.6 kbp) | 11,360 | 4179 (85.4%) | 2 (1-3) |
| MiDAS Global | 12,106 | 475.9 kbp (280.1 - 873.8 kbp) | 19,283 | 8825 (72.9%) | 1 (0-2) |
| Global Ocean Microbiome Catalogue | 10,180 | 14.4 kbp (8.9 - 29.7 kbp) | 11,336 | 5819 (57.2%) | 1 (0-2) |
| HumGut | 2,094 | 24.7 kbp (11.8 - 63.8 kbp) | 1,836 | 1106 (52.8%) | 1 (0-1) |

To validate the specificity of the newly constructed pHMMs, they were used to search for homologs in the GTDB. These were then clustered into BGCs using a maximum intergene distance of 5000 bp and compared to the homologs identified using the query genes. Bit score thresholds were set for each pHMM to improve their accuracy. Once this final step of epsProtocol was completed, an epsSMASH rule was made for each known exoPS using the newly made pHMMs (**Supplementary Note 1**). Because the sucrase-dependent systems only contain a single sucrase gene, we could not develop pHHMs and rules for these based on epsProtocol. We therefore built and tested pHMMs for fructansucrases and alpha-glucansucrases outside of the epsProtocol workflow (**Supplementary Note 1**).

epsSMASH performance was evaluated by comparing its predictive ability to previous attempts at predicting genomic potential for exoPS production (**Supplementary Note 2, Supplementary Figure 1-2**). This comparison showcased the accuracy of epsSMASH, as it detected true exoPS BGCs detected by previous studies, while avoiding the ones which upon manual inspection were deemed false positives. Additionally, epsSMASH detected 47 exoPS BGCs across 14 well-studied model organisms, 96% of which were deemed true positives based on literature evidence and manual validation (**Supplementary Table 2**).

### High-quality long-read genomes are required for identification of exoPS BGCs

Previous studies have shown that contiguity of the input genome data strongly influences antiSMASH results^17^. Long-read genome assemblies typically reveal more secondary metabolite BGCs per genome than short-read assemblies, which often miss or fragment repetitive regions^18^. Moreover, most BGCs identified in short-read assemblies are located near contig edges, suggesting that many are incomplete^19,20^. Although exoPS BGCs are not inherently repetitive, they usually contain many genes^5^, which could hinder complete assembly when using only short reads.

To assess the effects of different sequencing methods on the recovery of exoPS BGCs, we applied epsSMASH to three studies which assembled both long-read and short-read genome databases from the same samples (**Figure 2**): short- and long-read metagenome-assembled genomes (MAGs) from the Microflora Danica (MFD) Project^21,22^, short-, hybrid, and long-read MAGs from mouse fecal matter^19^, and short-read and hybrid MAGs from the Singapore Platinum Metagenomes Project (SPMP)^23^. The N50 values (defined as the contig length where the sum of all contigs of that size or longer is equal to at least half the genome size) in each study were higher in long-read, high quality (HQ) MAGs (MAGs with genome completeness >90% and contamination < 5%), compared to short-read, medium quality (MQ) MAGs (MAGs with genome completeness >50%), showing that sequence contiguity is indeed higher in HQ, long-read MAGs (**Figure 2**).

**Figure 2:**
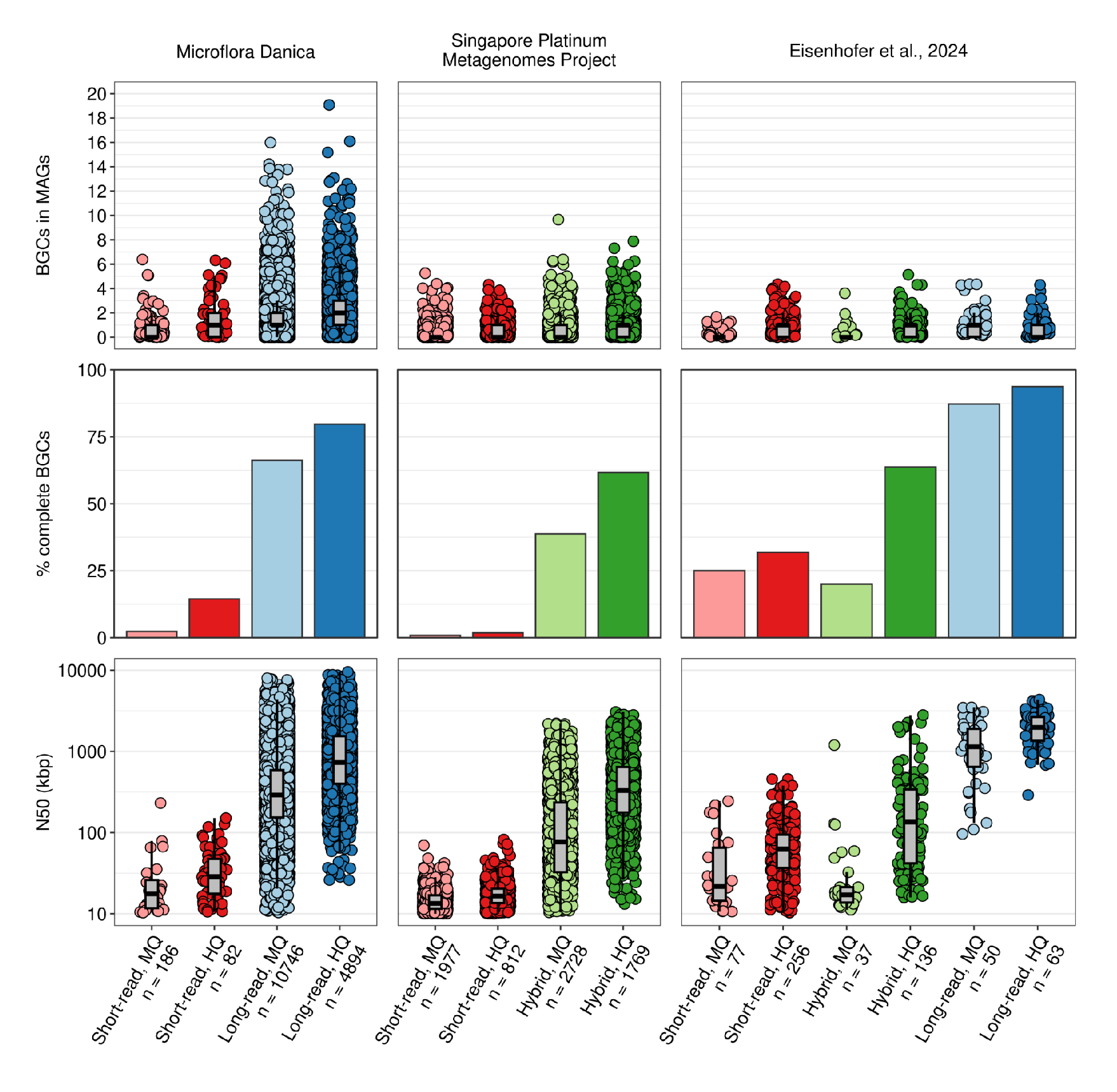
Benchmarking effects of contiguity on epsSMASH results. **Top:** Number of BGCs in the MAGS of three studies which used different sequencing methods on the same samples. **Middle:** Percentage of complete exoPS BGCs detected in the three studies. A BGC is designated incomplete if it is located near a contig edge. Colors indicate sequencing method and genome quality. Hybrid sequencing indicates that both long-read and short-read sequencing data were used to construct the MAGs. **Bottom:** Contig N50 values (kbp) for the MAGs in the three studies.

In the MFD and SPMP studies, long-read and hybrid genome assemblies result in more exoPS BGCs detected per MAG and HQ MAGs consistently result in more exoPS BGCs per MAG as compared to MQ MAGs. In the Eisenhofer genomes, epsSMASH detects a comparable amount of exoPS BGCs per MAG in HQ MAGs assembled from short-read, hybrid and long-read sequencing. In all three studies the percentage of complete exoPS BGCs (defined as BGCs not close to a contig edge) were highest in HQ MAGs assembled using long-reads or hybrid reads as compared to short-read assembly. These results show that high genome completeness and sequence contiguity is necessary if the goal is to comprehensively capture genomic potential for exoPS production in a microbiome.

### Most exoPS BGCs in environmental microbiomes are uncharacterised

epsSMASH provides a unique opportunity to examine the genetic potential for exoPS production in microbiomes where biofilm formation and composition are of ecological or technical relevance. We therefore examined four major ecosystems, the human gut, soil, ocean and activated sludge from wastewater treatment plants. For each environment, we prioritised HQ MAG catalogues assembled with long-read sequencing which covered a large geographic area.

In the MFD project, a genome catalogue was assembled from deep, long-read sequencing of 154 soil and sediment samples collected across Denmark^21^. The MiDAS Global genome catalogue is a collection of MAGs constructed from deep long-read sequencing of 83 activated sludge samples from wastewater treatment plants across the world (Manuscript in preparation). The Global Ocean Microbiome Catalogue (GOMC) is a collection of genomes constructed using a globally distributed set of ocean metagenomes and marine microbial genomes obtained from a selection of public databases^24^. The HumGut catalogue was constructed by screening over 5,700 healthy human gut metagenomes for the presence of genomes from RefSeq and the Unified Human Gastrointestinal Genome collection^25^. Median contig N50 values were an order of magnitude higher in the MFD and MiDAS Global catalogues compared to GOMC and HumGut (**Table 1**), likely due to the abundance of short-read sequences in public genome databases^26^.

We ran epsSMASH on the HQ species-representative genomes of each catalogue (**Supplementary Table 3**) and found exoPS BGCs in 52.8-85.4% of the MAGs, with a median of 1-2 exoPS BGCs per MAG (**Table 1**). The percentage of complete exoPS BGCs in the MFD and MiDAS Global catalogues were 79.7% and 85.9%, while the percentages in GOMC and HumGut were around 50% (**Figure 3a**). The higher levels of exoPS BGC detection and higher percentages of exoPS BGC completeness in the MFD and MiDAS Global catalogues are likely due to their higher sequence contiguity (**Table 1**). We found extreme outliers in all four databases (**Figure 3b**). In members of the Cyanobacteriales order we found species with up to 19 distinct exoPS BGCs, with genomes in the order containing on average 4.2 BGCs/MAG (**Supplementary Table 3**). Cyanobacterial exoPS production has been studied extensively, but few studies have focused on functional properties^27^. The Cyanobacterial genome which contained 19 exoPS BGCs belonged to the genus *Rivularia*, which are known to produce thick mucilaginous sheaths composed of acidic polysaccharides^28^.

**Figure 3:**
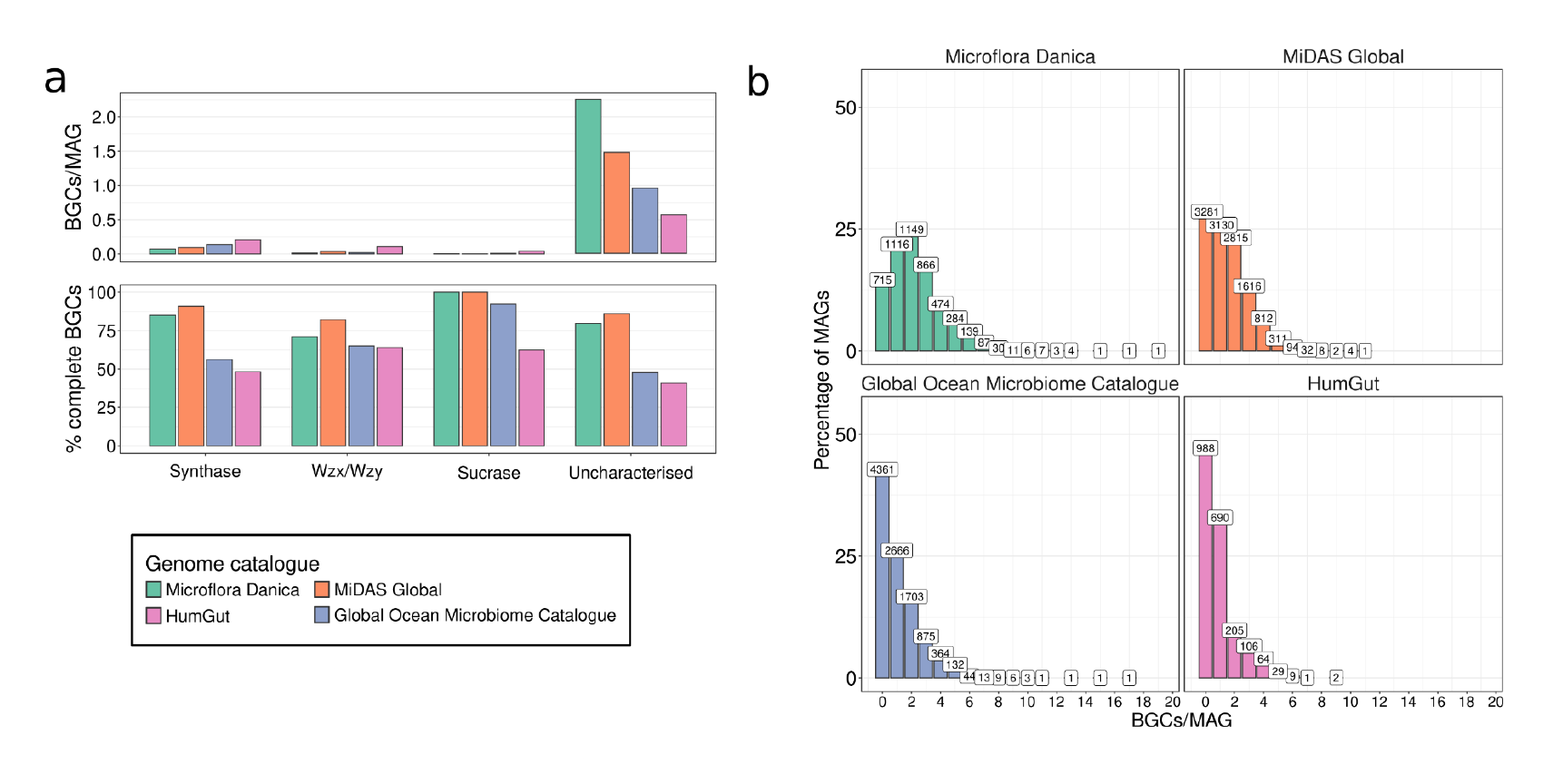
epsSMASH results for the four genome catalogues. **a**. Average number of uncharacterised, synthase-, Wzx/Wzy and Sucrase-dependent exoPS BGCs detected by epsSMASH per MAG and percentages of how many of these are complete in the four genome catalogues. A BGC is designated incomplete if it is located near a contig edge. **b**. Percentage of MAGs in each genome catalogue containing different numbers of BGCs/MAG.

In all four genome catalogues, the majority of exoPS BGCs (62%-96%) were detected by loose rules, meaning that they belong to uncharacterised Wzx/Wzy- or ABC transporter-dependent exoPS pathways (**Figure 3b**). When looking only at exoPS BGCs detected by strict and relaxed rules, HumGut contained the most exoPS BGCs/MAGs. This is likely because the human gut represents the most well studied and least diverse environments of the four analysed in this study.

### exoPS potential in core activated sludge genera

To gain insight into the exoPS potential of the most prevalent bacteria in global activated sludge plants, we analysed the exoPS BGCs detected in the “strict core genera”, defined by Dueholm et al. (2022) as genera with >0.1% abundance in more than 80% of activated sludge wastewater treatment plants worldwide (**Figure 4**). In every strict genus but *Bdellovibrio*, the vast majority of species encode at least one exoPS BGC. In several genera (*Pirellula, Terrimonas, Ferruginibacter* and *Flavobacterium*) all exoPS BGCs belong to uncharacterised exoPS pathways. Genera from the Burkholderiales order (*Sulfuritalea, Zoogloea, Azonexus*, midas_g_81, *Acidovorax* and *Lautropia*) contain species with Pel, zooglan and poly-N-acetylglucosamine (PNAG) BGCs. *Rhodobacter* contains species with alginate BGCs, while *Novosphingobium, Haliangium, Azonexus* and *Acidovorax* are the only strict core genera with species encoding cellulose BGCs. More than half of the species in the *Hyphomicrobium* genus encode a pel-like BGC, indicating BGCs which lack the complete pel operon, but contain the biosynthetic core genes *pelF* and *pelG*. Large interspecies differences can be seen for the exoPS BGCs in the strict core AS genera, with anywhere from 1%-86% of species in a genus encoding a given exoPS BGC. The observation that exoPS BGCs can be phylogenetically widespread, yet inconsistently conserved at a given taxonomical level is corroborated by previous analyses of exoPS BGCs, and has also been observed for other types of BGCs^10–12^.

**Figure 4:**
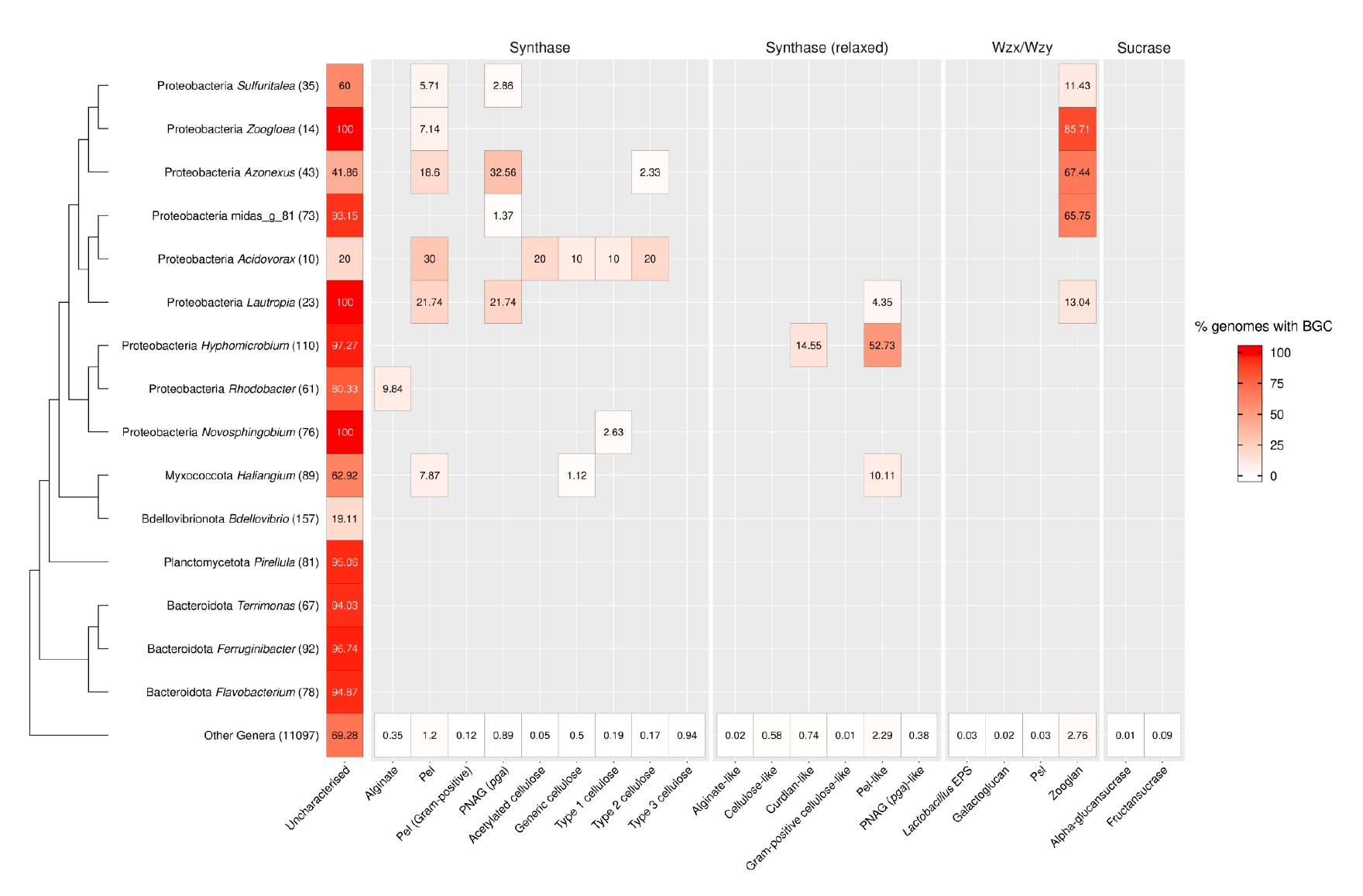
exoPS BGCs detected in core activated sludge genera. exoPS BGCs encoded by strict core activated sludge genera from the MiDAS Global genome catalogue. Phylum and genus names are based on MiDAS 5.3 taxonomy. Numbers next to genus names indicate the number of species present in MiDAS Global from that genus. Numbers inside heatmap tiles indicate the percentage of species within a given genus that encodes an exoPS BGC.

This phenomenon may be explained in part by the “selfish operon model”^30^, which proposes that clustering functionally related genes promotes their horizontal transfer, thereby increasing the likelihood that the gene cluster (and its constituent genes) successfully colonise new taxa.

### Gene cluster families reveal novel and phylogenetically widespread exoPS BGCs

epsSMASH is based on the antiSMASH platform, enabling epsSMASH users to take advantage of the bioinformatic software built specifically for antiSMASH output, such as BiG-SCAPE^31^.

BiG-SCAPE identifies gene cluster families (GCFs) by expressing the differences in Pfam domain similarity, sequence similarity and synteny as distances between BGCs. GCFs are assigned by BiG-SCAPE by clustering BGCs with distances below a user-defined cut-off. We used BiG-SCAPE to create gene cluster similarity networks of the exoPS BGCs from each genome catalogue (**Figure 5**). Since each genome catalogue only contains species representatives, the largest GCFs represent exoPS BGCs that are present in the highest number of distinct species. Consequently, these GCFs contain BGCs with broad taxonomic representation, making them a suitable starting point for identifying both widespread and potentially novel exoPS BGCs. The GCFs were sorted by number of BGCs, and the 20 largest GCFs were selected for further analysis.

**Figure 5:**
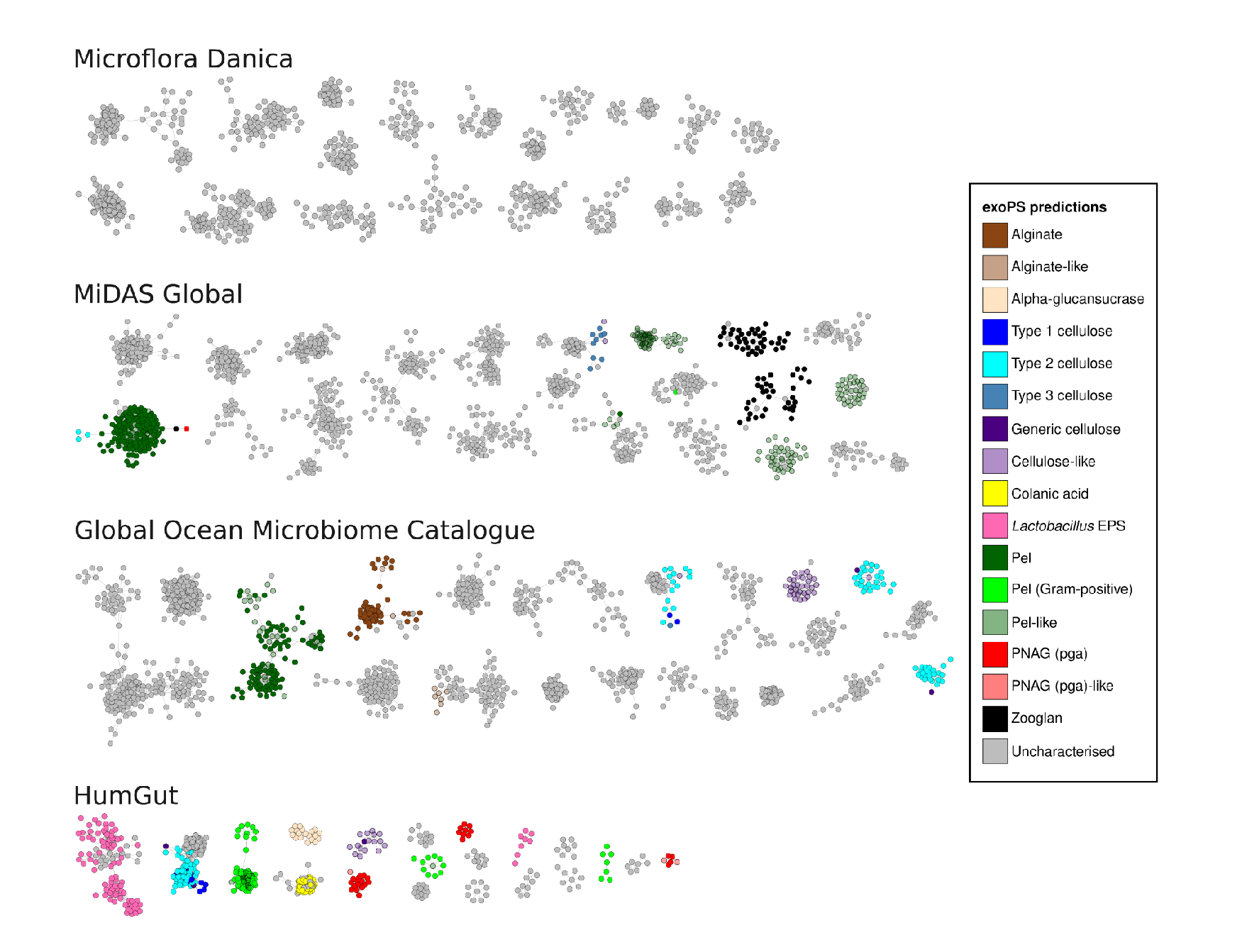
Gene cluster similarity networks. The 20 clusters with the most exoPS BGCs in the gene cluster similarity networks constructed for each genome catalogue. BGCs are colored by the epsSMASH rule they were detected by. BGCs detected by relaxed rules (i.e. those with -like suffix) are colored the same as their strict rule counterparts with 90% opacity. BGCs detected by loose rules are considered uncharacterised.

In all four gene cluster similarity networks, exoPS BGCs detected by the strict rules are grouped into distinct GCFs. Occasionally, exoPS BGCs detected by a single strict rule form several distinct GCFs, representing different taxonomic groups encoding the same BGC (**Supplementary Figure 3-6**). GCFs representing an exoPS BGC matching a strict rule often also contain exoPS BGCs detected by relaxed or loose rules. These invariably represent truncated versions of the exoPS BGCs detected by the strict rules. Cases like these highlight the ability of relaxed rules to catch synthase-dependent systems that are missed by their strict rule counterparts, enabling exoPS BGC detection even in fragmented genomes or near contig edges.

In every gene cluster similarity network but Microflora Danica, one of the three largest GCFs consist of pel and pel-like BGCs, indicating that pel is a widespread exoPS in several ecosystems. Remarkably, epsSMASH detected pel BGCs in 14 distinct phyla, making it the most phylogenetically widespread exoPS BGC in the human gut, ocean and activated sludge microbiomes. Further analysis that rearrangements in pel operon synteny across phyla appeared constrained by the protein–protein complexes formed within the operon (**Supplementary Note 3, Supplementary Figure 7**).

The MiDAS Global network contains three large GCFs composed entirely of “pel-like” BGCs. These represent widespread putative exoPS BGCs which contain homologues of the strongly conserved *pelFG* genes, but where the complete pel operon is not detected. Until now, pel operons had not been discovered in the taxonomic classes containing these BGCs. Using structural homology searches we showed that these BGCs were indeed distant homologs of the canonical *pelABCDEFG* operon (**Supplementary Table 4**). We constructed additional pHMMs for the pel epsSMASH rule from these distantly homologous sequences, improving the annotation of pel operons in epsSMASH (**Supplementary Note 4, Supplementary Figure 8**). This example demonstrates how relaxed rules can be used to detect distant homologs of known systems by targeting only their conserved core genes.

In every gene cluster similarity network but HumGut, the majority of the 20 largest GCFs are composed solely of uncharacterised exoPS BGCs (**Figure 5**). This was especially true for the MFD catalogue, where the 20 largest GCFs were all composed of uncharacterised exoPS BGCs. GCFs of both known and uncharacterised exoPS BGCs clustered phylogenetically, often at the order, family or even genus level (**Supplementary Figure 3-6**). We suggest that each uncharacterised GCF represents a novel, phylogenetically conserved type of exoPS BGC, and that the most widespread of these clusters could be important EPS components in their respective environments.

### A Platform for Exploring and Characterizing New exoPS BGCs

To begin exploring the uncharacterised exoPS BGCs in environmental communities, we focused on the largest GCF in the MiDAS Global gene cluster similarity network (**Figure 5**). The exoPS BGCs in this GCF all belong to genera from the Sphingomonadaceae family. Several genera from this family (*Sphingobium, Sphingopyxis, Novosphingobium*) are part of the core microbiome in activated sludge^29^ (**Supplementary Figure 9**). The Sphingomonadaceae family has previously been shown to contain exoPS producers such as the genus *Sphingomonas* which produce sphingans and promonan^32,33^ as well as sanxan, which is produced by several genera within the family^34^. However there is neither syntenic nor sequence similarity between this novel exoPS BGC and previously characterised exoPS pathways in Sphingomonadaceae. We created a new epsSMASH rule, “sphingomonadales_eps”, to describe the novel BGC using genes from the model organism *Sphingopyxis granuli*. We then ran epsSMASH on all high quality, species representative genomes from the Sphingomonadales order in GTDB v220 (n=902), revealing that almost all members of the order possess this exoPS BGC, and that the synteny is remarkably conserved (**Figure 6a,b**).

**Figure 6:**
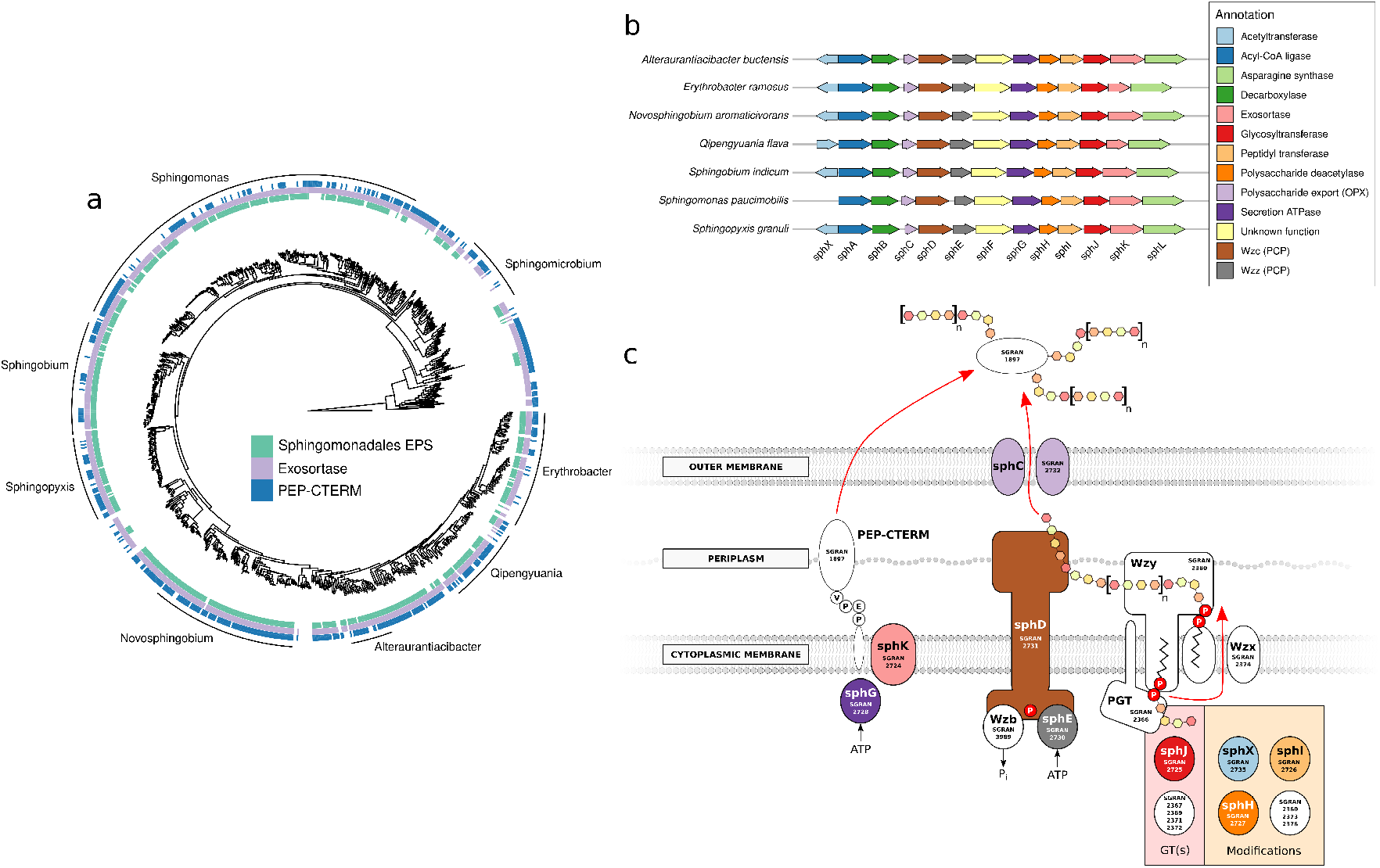
Analysis of a novel Sphingomonadales exoPS BGC. **a**. Phylogenomic tree of the 902 high quality species representative Sphingomonadales genomes in the Genome Taxonomy Database v220. The inner ring indicates each genome where the Sphingomonadales exoPS BGC is detected. The middle ring indicates each genome that contains an exosortase homologue (TIGR02602). The outer ring indicates each genome that contains at least one PEP-CTERM homologue (TIGR02595). The tree was created using gtdb-tk v2.5.1. **b**. Gene arrow plots of the Sphingomonadales exoPS BGC from members of the 7 most populated genera in the Sphingomonadales genome set (Excluding Sphingomicrobium which does not contain the BGC). **c**. Proposed biosynthetic pathway of the Sphingomonadales exoPS BGC. A phosphoglyceryltransferase (PGT) primes the undecaprenyl phosphate with an intitial sugar monomer. One or more glycosyltransferases (GTs) add more sugar monomers to the repeating unit, while other enzymes (e.g. acetyltransferases, acyltransferases, deacetylases) modify the sugar monomers. Repeat units are flipped into the periplasm by Wzx, whereafter they are polymerised by Wzy. A polysaccharide co-polymerase (PCP) determines the extent of the polymerisation and guides the nascent polysaccharide to the outer membrane polysaccharide export (OPX) protein, which transports them outside the cell. The activity of the PCP protein is regulated by phosphorylation and dephosphorylation, which is mediated by a tyrosine kinase (Wzz) and a tyrosine phosphatase, respectively. The exosortase recognises PEP-CTERM bound to the cytoplasmic membrane and cleaves them, releasing them into the periplasm after which they eventually are secreted into the extracellular space. Here, they are glycosylated by the polysaccharides secreted through the OPX. The colors of the proteins indicate which gene in the Sphingomonadales BGC they are encoded by (see **b**). White genes were found in other loci in the *S. granuli* TFA genome, as indicated by the SGRAN gene ids.

When running epsSMASH, an optional setting called Clusterblast can be enabled, which compares detected exoPS BGCs with a database containing the BGCs which were manually validated in epsProtocol (**Figure 1a**). The Sphingomonadales exoPS BGCs shared remarkable synteny and sequence similarity to part of the zooglan BGC, another exoPS from Pseudomonadota which is also widespread in activated sludge genomes^35^. This connection as well as the presence of an *xrtA* exosortase homologue suggests that this exoPS BGC is involved in the PEP-CTERM/exosortase protein sorting system^36^. This system, which is found in many biofilm-producing bacteria from environmental sources^36^, facilitates the export of proteins containing Pro-Glu-Pro (PEP) motifs into the periplasm from which they are thought to be incorporated into the extracellular polysaccharide layer^37^. In *Zoogloea*, floc formation is only possible when both zooglan genes and the PEP-CTERM/exosortase system are expressed^37^.

We performed a search of the 902 Sphingomonadales genomes using the TIGRFAM pHMMs for the *xrtA* exosortase (TIGR02602) and the PEP-CTERM domain (TIGR02595) (**Figure 6a**). PEP-CTERM genes, which have been described as “phylogenetically sporadic and somewhat sparse”^36^, were shown to be widespread in the Sphingomonadales order. In line with previous studies, PEP-CTERM-positive genomes almost invariably contained an exosortase as well. The reverse was not true, however, as many exosortase-containing genomes did not contain homologs to the PEP-CTERM pHMM. Interestingly, many Sphingomonadales genomes without the newly discovered Sphingomonadales exoPS BGCs still contained the exosortase/PEP-CTERM system. We suggest that these systems depend on other exoPS BGCs, since epsSMASH detected at least one uncharacterised exoPS BGC in every Sphingomonadales genome. We hypothesise that this newly discovered exoPS BGC performs an analogous function to the PEP-CTERM/exosortase-associated zooglan system, and propose a possible biosynthetic pathway for the system based on the genes found in *Sphingopyxis granuli* TFA (**Figure 6c**). Further phylogenetic analysis of this exoPS BGC is needed to uncover the origins and diversity of this system, and laboratory-based methods (mutagenesis, physical characterisation of exoPS structure) must be carried out to fully characterise this novel exoPS. Nevertheless, this analysis showcases the usefulness of epsSMASH for exploring the vast number of uncharacterised exoPS systems.

## Conclusions

We developed epsSMASH, the most comprehensive high-throughput tool for characterising the genomic potential for exoPS production so far. epsSMASH is available both as a user-friendly web service for single genome analyses and a command-line tool for large-scale analysis of genome-resolved metagenomes. When compared to previous attempts at predicting exoPS potential, epsSMASH was able to detect genuine exoPS BGCs while avoiding false positives. Applying epsSMASH to four large genome databases representing both environmental and human microbiomes led to the detection of exoPS BGCs in 52.8-85.4% of the genomes. exoPS BGC recovery and completeness was highly dependent on the level of contiguity of the analysed genomes. While a median of 1-2 BGCs per MAG were detected, some species contained up to 19 exoPS BGCs per MAG. Pel was the most phylogenetically widespread exoPS BGC in the human gut, ocean and activated sludge microbiomes, but was rarely detected in bacterial genomes from soil. Most bacterial exoPS BGCs in all four environments were uncharacterised. Constructing GCFs from the epsSMASH output allowed us to identify and define widespread, previously uncharacterised systems, providing a solid foundation for downstream experimental characterisation. We anticipate that epsSMASH will become an essential tool for investigating exoPS production in microbial communities, leading to an improved understanding of the components in the ubiquitous and extremely dynamic biofilm matrix.

## Methods

### epsSMASH predictions

Genome fasta files containing nucleotide sequences (contigs) were used as input for epsSMASH (v1.0) with the “--genefinding-tool prodigal” flag. The snakemake workflow Multismash (v0.5.2)^38^, was minimally modified to use epsSMASH instead of antiSMASH and was used for parallelising the epsSMASH runs and for generating overview files for the epsSMASH results. Gene-level information was extracted from Multismash results using a custom python script (https://github.com/AOHD/multismash_parser).

### Gene cluster families

From the epsSMASH results for each genome catalogue, we created gene cluster similarity networks with BiG-SCAPE 2.0^31^, using the cluster command with the --force-gbk flag and a Gene Cluster Family (GCF) cut-off value of 0.4.

### Data visualisation

GCFs were annotated and arranged using Cytoscape (v3.10.3)^39^. All other figures were generated in R (v4.4.0) using the following packages: gggenes (v0.5.0)^40^, tidyverse (v2.0.0)^41^, data.table (v1.16.4)^42^, ggplot2 (v3.5.2)^43^, ggtree (v3.14), ggtreeExtra (v3.14) and tidytree (v0.4.6)^44^. All figures were processed in Inkscape v1.1.2.

### Phylogenomic trees

GTDB-tk (v2.4.1)^45^ was used to create a multiple sequence alignment (MSA) of the phylogenetic marker genes in the 902 HQ Sphingomonadales genomes from GTDB (v220)^16^. Iqtree2 (v2.2.6)^46^ was used to construct a phylogenomic tree from this MSA, using two genomes from the Rickettsiales as an outgroup (Outgroup accessions: RS_GCF_021378375.1 and RS_GCF_000215485.1). A multiple sequence alignment of the phylogenetic marker genes was created from MiDAS 4 strict core.

### Code availability

The epsSMASH command-line tool can be found at https://github.com/AOHD/epsSMASH, while the web-service is available at http://epssmash.org/. Documentation for the tool is available at https://docs.epssmash.secondarymetabolites.org/. Code used to generate and visualise the results in this article is available at https://github.com/AOHD/epsSMASH_paper.

## Acknowledgements

The study was funded by the Novo Nordisk Foundation (Grant NNF22OC0071498)

## Notes

https://github.com/AOHD/epsSMASH

http://epssmash.org/

https://docs.epssmash.secondarymetabolites.org/

https://github.com/AOHD/epsSMASH_paper

https://github.com/cmc-aau/epsProtocol

### Competing Interest Statement

The authors have declared no competing interest.

