## Supplemental Information for "epsSMASH uncovers exopolysaccharide biosynthetic gene clusters in environmental and human microbiomes"

**Table of contents:**

Supplementary Figure 1-8
Supplementary Note 1-4
Supplementary References

### Supplementary Figures

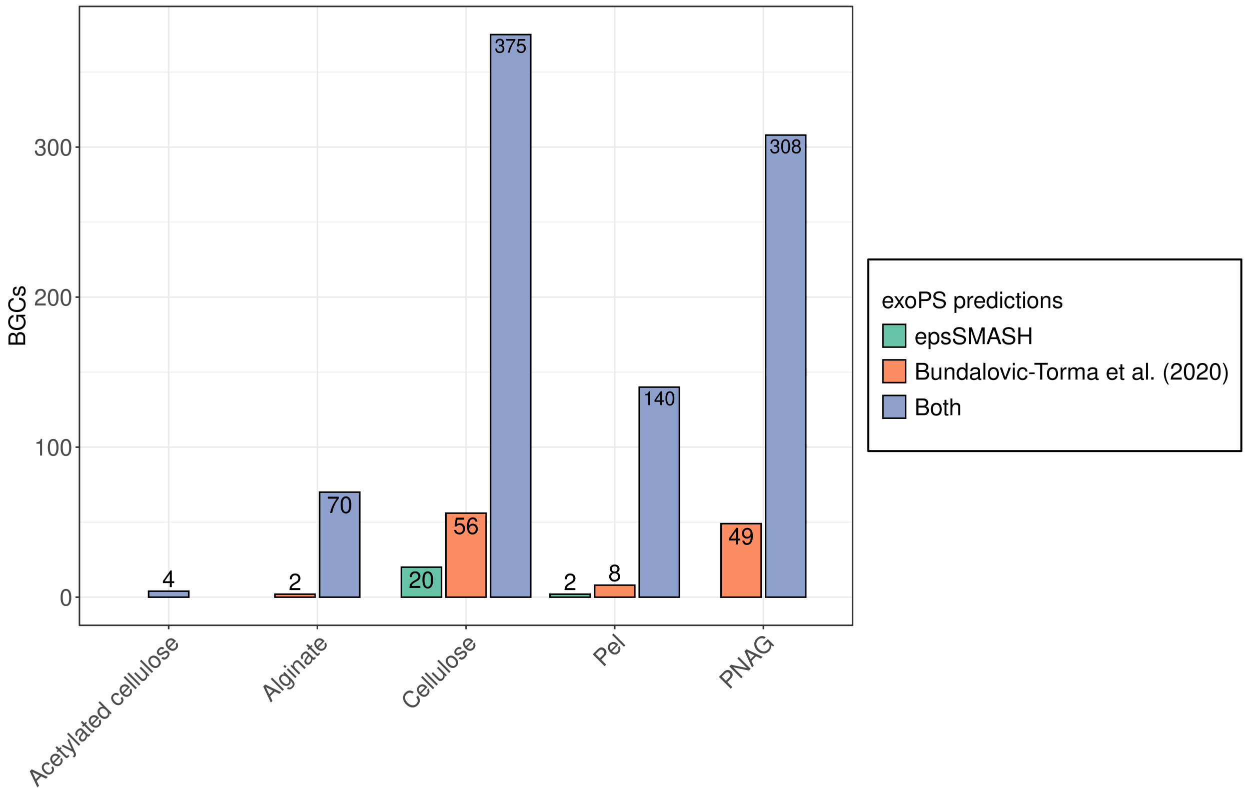


**Supplementary Figure 1:** exoPS BGCs detected by epsSMASH, Bundalovic-Torma et al. (2020) or both in bacterial genomes obtained from NCBI (2015).

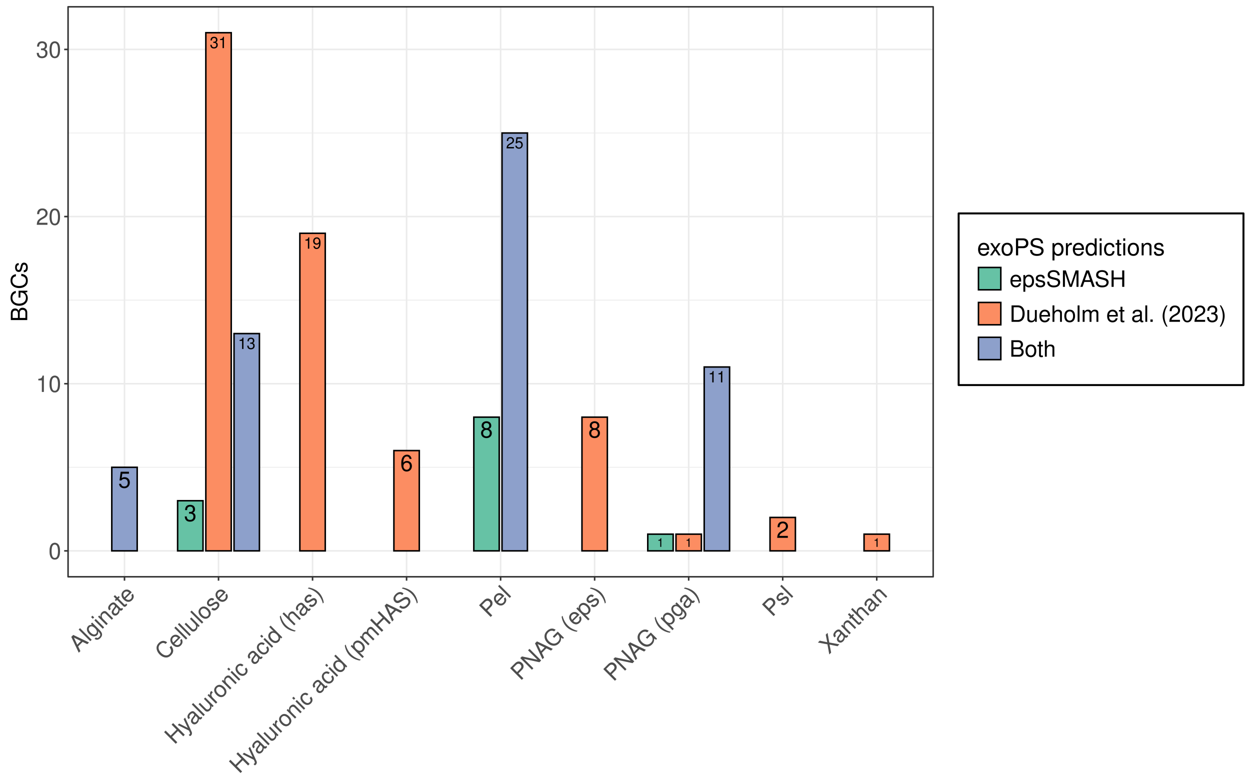


**Supplementary Figure 2:** exoPS BGCs detected by epsSMASH, Dueholm et al. (2023) or both in MAGs obtained from Danish activated sludge wastewater treatment plants.

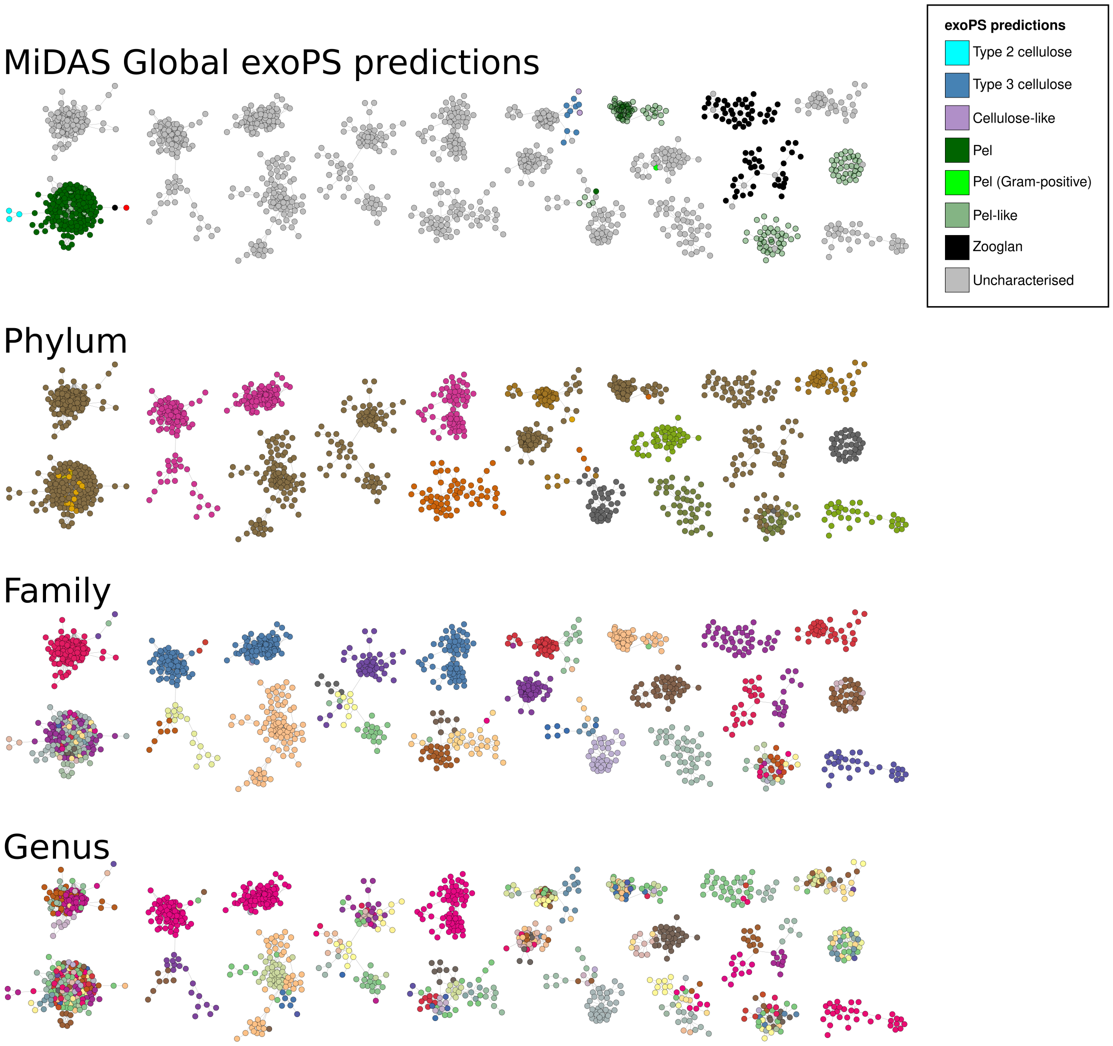


**Supplementary Figure 3:** 20 largest GCFs in the MiDAS Global gene cluster similarity network. The topmost figure shows the GCFs colored by exoPS predictions, while the following three figures are colored by phylum, family and genus, respectively.

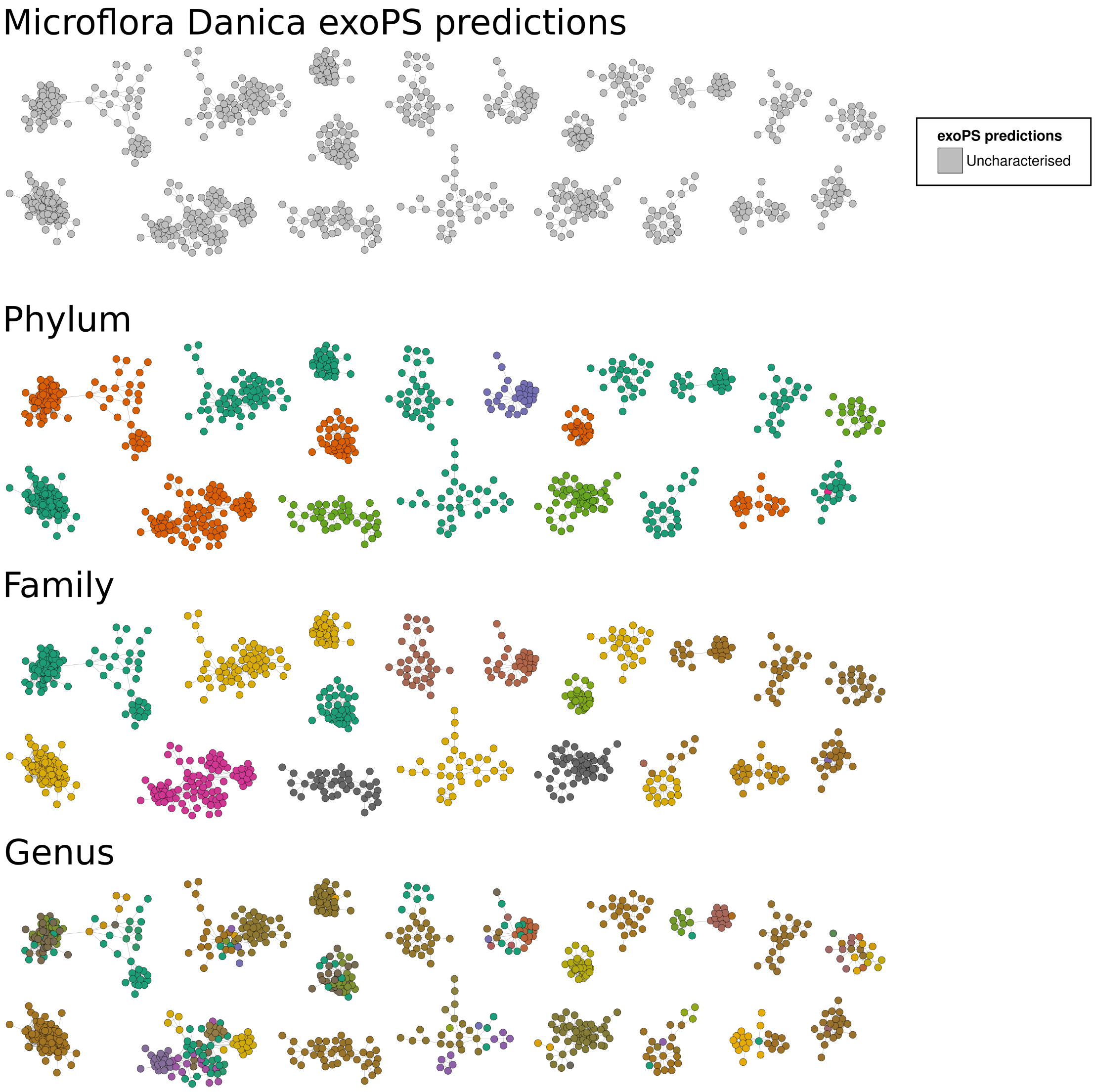


**Supplementary Figure 4:** 20 largest GCFs in the Microflora Danica gene cluster similarity network. The topmost figure shows the GCFs colored by exoPS predictions, while the following three figures are colored by phylum, family and genus, respectively.

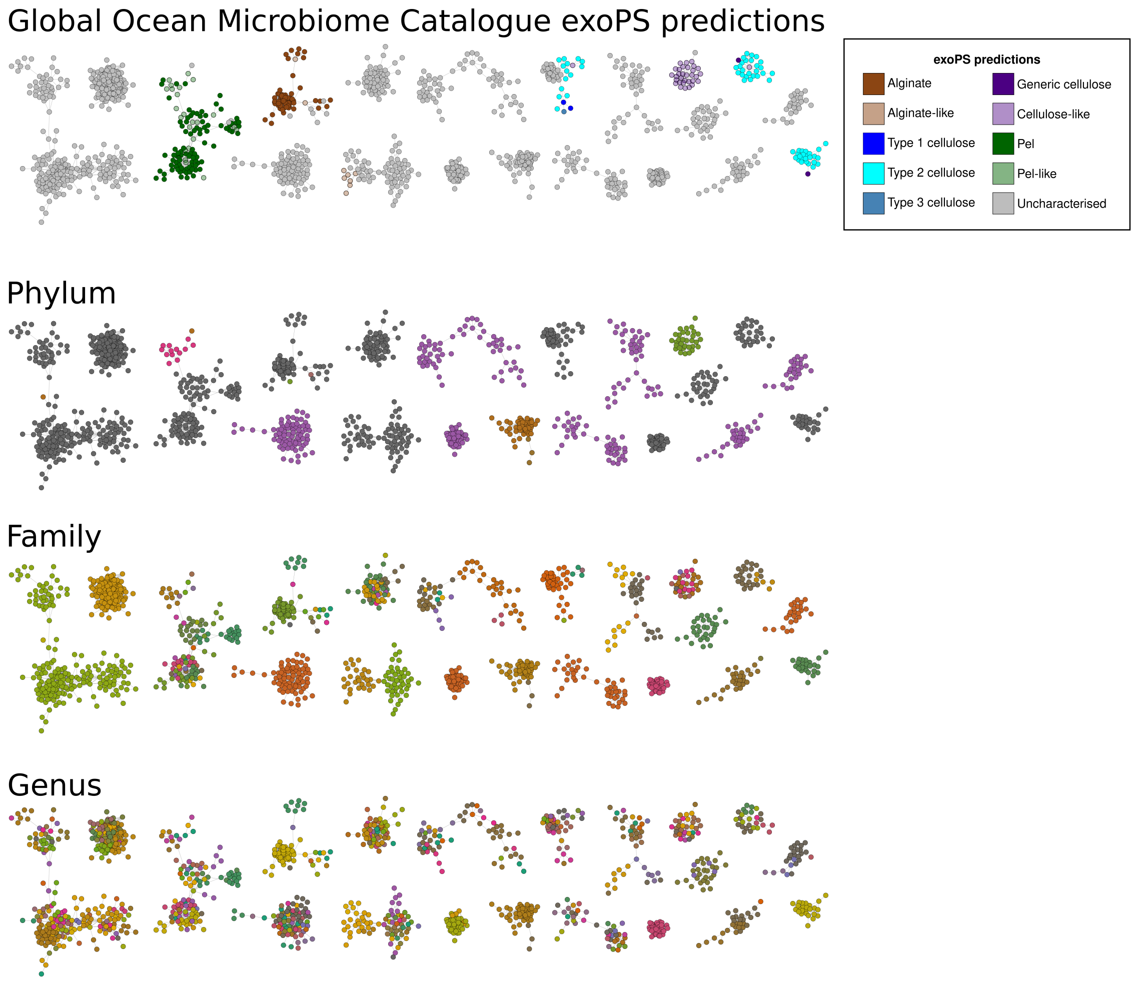


**Supplementary Figure 5:** 20 largest GCFs in the Global Ocean Microbiome Catalogue gene cluster similarity network. The topmost figure shows the GCFs colored by exoPS predictions, while the following three figures are colored by phylum, family and genus, respectively.

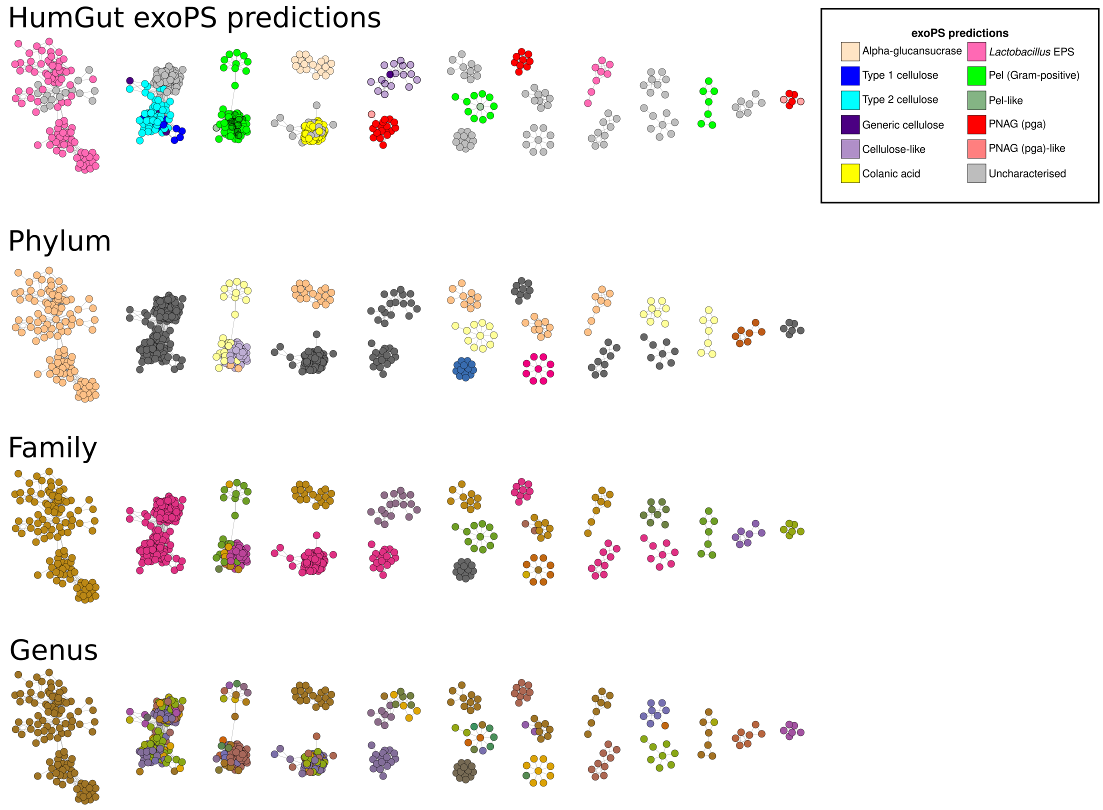


**Supplementary Figure 6:** 20 largest GCFs in the HumGut cluster similarity network. The topmost figure shows the GCFs colored by exoPS predictions, while the following three figures are colored by phylum, family and genus, respectively.

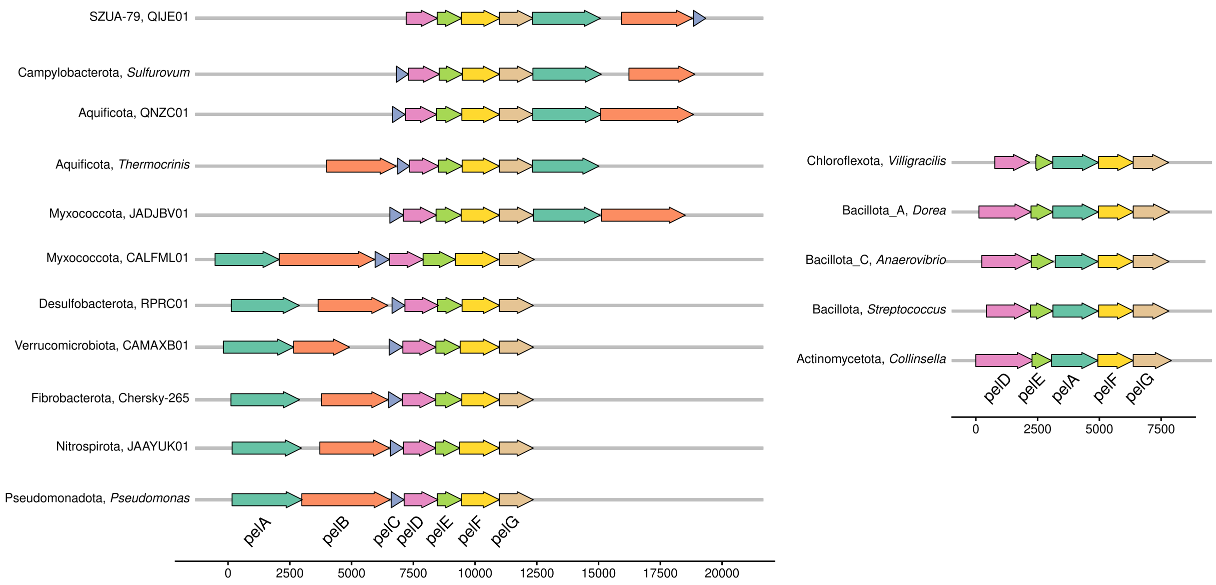


**Supplementary Figure 7:** Representative pel operons from each phylum in which epsSMASH detected pel in the four genome catalogues. Phylum and genera (GTDB taxonomy) are listed next to the operons.

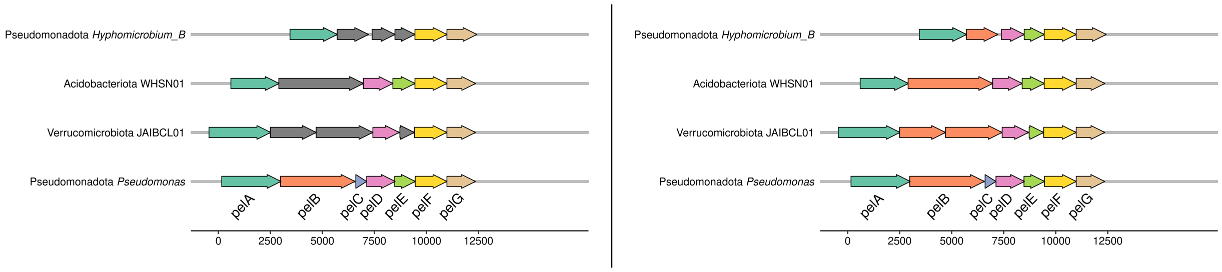


**Supplementary Figure 8: Left:** Representative operons from the three "pel-like" BiG-SCAPE clusters in the MiDAS sequence similarity network, aligned to pelG of the *Pseudomonas aeruginosa* pel operon. Phylum and genera (GTDB taxonomy) are listed next to the operons. **Right:** The same operons after structural homology search annotation of unknown genes with Foldseek.

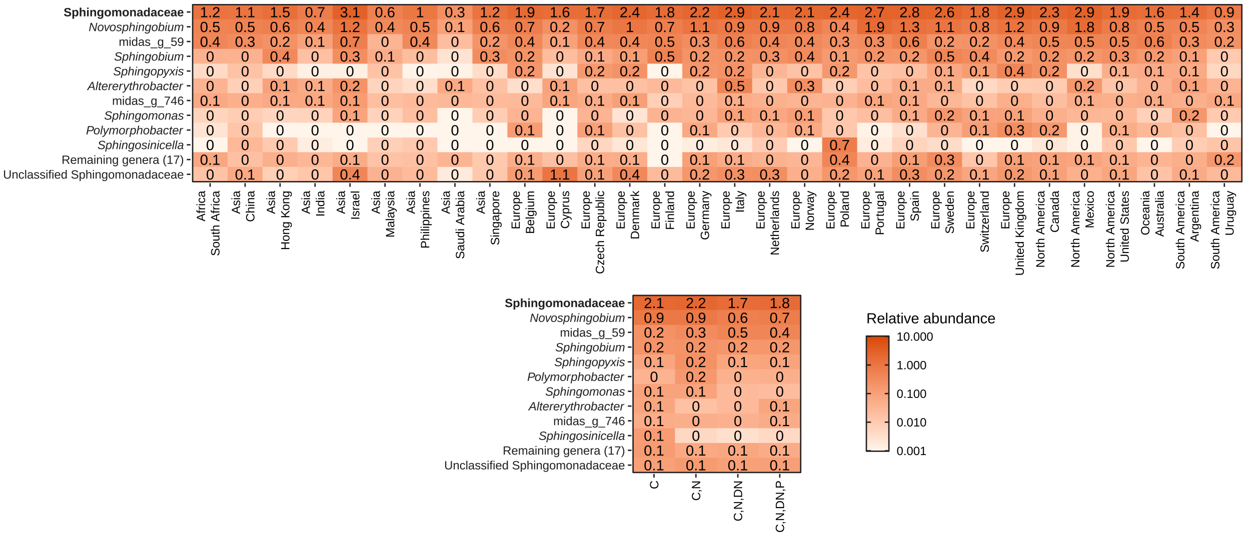


**Supplementary Figure 9:** Relative abundances of Sphingomonadaceae genera in activated sludge across countries (top) and different wastewater treatment process types (bottom). The top row shows the abundances of the Sphingomonadaceae family as a whole. Based on V1-V3 amplicon data from MiDAS 4 (Dueholm et al., 2022).

**Supplementary Note 1: Generation of pHMMs and rules for BGC prediction: epsProtocol**

A reproducible Snakemake workflow, epsProtocol, was developed to generate pHMMs and BGC prediction rules for epsSMASH (**Figure 1b**). A total of 21 known and characterised exoPS BGCs covering three of the four known pathways for exoPS biosynthesis (the Wzx/Wzy-, ABC-transporter and Synthase-dependent pathways, see **Supplementary Table 1**) were chosen. Each of these BGCs were subjected to the epsProtocol workflow, resulting in at least one epsSMASH rule. Code for the epsProtocol workflow can be found at https://github.com/cmc-aau/epsProtocol, including knitted Rmarkdown files for each rule created using the workflow.

**Gathering known and characterised exoPS BGCs**

The epsProtocol workflow builds on an extensive literature review of the exoPS BGC of interest. The information about each gene in the gene cluster, the observed synteny/syntenies of the BGC, which taxonomic groups are known to harbor the BGC as well as information about the structure and function of the excreted exoPS was summarised (**Supplementary Table 1**). In cases where exoPS production was described in organisms across different genera, we selected one representative from each, resulting in more than one amino acid sequence for each gene. Finally, each gene was assigned one of six functions, based on knowledge gained from the literature review: regulation (genes which regulate the pathway), modification (genes which modify the exoPS, e.g. acetyltransferases), degradation (genes which cleave the exoPS), transportation (genes which facilitate exoPS secretion), polymerisation (genes which contribute sugar monomers to the growing exoPS, e.g. glycosyltransferases) and polymerisation/transportation (genes which both add sugar monomers to the growing exoPS and facilitate exoPS secretion, e.g. synthases and the Wzy polymerase) (**Supplementary Table 1**).

**BGCs manual curation**

We looked for sequence homologs of the genes in the genomes of the 113.106 species representatives in the Genome Taxonomy Taxonomy Database (GTDB) (v220)[1] using the iterative protein search tool Jackhmmer (HMMER v3.1)[2] with an e-value threshold of 0.1 (-E 0.1). This permissive search was an attempt to capture all amino acid sequences with even the slightest sequence similarity to our representative exoPS BGC genes in the GTDB. The protein hits from the Jackhmmer search were grouped into putative BGCs using an in-house R-script which clusters genes with an inter-gene distance below 5000 bp.

The putative exoPS BGCs were manually curated using a custom R Markdown script. The primary goal of this step was to refine the search results, keeping only BGCs with gene composition and synteny matching the gene clusters described in literature, and discarding false positives. The first refinement of the putative BGCs consisted of designating one or more "core genes", removing all BGCs which did not contain these. Core genes were chosen based on existing literature and general knowledge about exoPS biosynthesis pathways, usually consisting of genes related to the transport and polymerisation of the exoPS. A minimum number of genes was set for the gene clusters;  determined by constructing histograms of gene distributions for different minimum gene requirements. A satisfactory minimum gene requirement would show a histogram with an even distribution of genes in the BGCs, indicating that one instance of each gene is present in the detected gene clusters. In addition to core gene presence and minimum gene count per BGC, we applied a third filter based on synteny patterns described in literature. Each putative BGC which contained all core genes and passed the minimum gene requirement were manually validated. This step allowed us to adjust the core gene set and minimum gene thresholds as needed. Our aim was to only keep the BGCs which had the synteny and core gene composition of the exoPS BGC described in literature.

The result after the manual validation of each putative exoPS BGC was a collection of exoPS BGCs which matched the gene synteny and composition of the exoPS BGC of interest. To enable comparison of novel exoPS BGCs detected by epsSMASH with manually validated systems, these collections were used to create the ClusterBlast database (**Figure 1a**).

**Constructing pHMMs from manually validated exoPS BGCs**
The manually validated BGC collection was used to construct pHMMs for each gene in the BGC, which were then used to construct epsSMASH detection rules. For all genes in the BGC collection the amino acid sequences were extracted and dereplicated at 90% average nucleotide identity using the usearch -clusterfast command with the -sort length and -centroids flags (v11.0.667) (Edgar, 2010). Each set of dereplicated genes was then aligned with MAFFT (v7.525)[3] using the einsi method, creating a multiple sequence alignment (MSA) for each gene in the BGC. These BGCs were first trimmed by TrimAl (v1.4.1)[4] using the -gt 0.75 flag to remove any columns with >25% gaps and then with a custom python script to remove any sequence where >25% of positions are gaps. To make sure that these trimmed MSAs did not contain any phylogenetic outliers, phylogenetic trees for each MSA were created using iqtree2 (v2.2.6)[5] with the flags -m WAG+R and -bb 1000. If any outliers were found, the BGC they belonged to was inspected, discarding either the gene or the entire BGC from the manually validated BGC collection. Once no outliers remained to be inspected, dereplication, alignment and trimming was performed again on the BGC collections. Profile HMMs for each gene in the BGC were then constructed using hmmbuild (HMMER v3.1)[2].

**Testing pHMMs and creating epsSMASH rules**
We performed a final validation to ensure that our pHMMs were accurately targeting the desired gene clusters. A homology search on all species representatives in the GTDB (v220)[1] with the hmmsearch tool (HMMER v3.1)[2] and an e-value threshold of 0.1 (-E 0.1) was performed with the pHMMs as queries. The protein hits were clustered into putative exoPS BGCs using the same in-house R-script used for the Jackhmmer results.

The putative exoPS BGCs obtained from the hmmsearch and subsequent gene clustering were filtered using the core gene sets and minimum gene requirements that were set during the curation of the manually validated exoPS BGC collection. The BGCs which passed this filtration were then compared to the manually validated BGC collection which the pHMMs had been created from. The majority of the exoPS BGCs detected by the pHMMs were also part of the manually curated BGC collection and were considered "true positives", as we were confident that the manually validated BGCs were genuine. BGCs from the manually validated collection which were not in the pHMM search results were considered false negatives for the same reason.

Each pHMM used in epsSMASH requires a bit score threshold. Bit scores were assigned to allow all hits which were present in the manually validated BGC collection, while removing any new hits which were deemed false positives based on manual inspection. After curating bitscores for each pHMM in the exoPS BGC, an epsSMASH rule was created using the designated bitscores and the chosen core/minimum gene requirements.

Most characterised exoPS BGC resulted in a single epsSMASH rule. The synthase-dependent *Bcs* system, encoding the biosynthesis pathway for Gram-negative cellulose, did not adhere to this. The Gram-negative cellulose biosynthesis pathway is a well-studied system, and known to produce different types of cellulose depending on the gene content in the *Bcs* operon. Leveraging this knowledge prompted us to create specific rules for four different types of cellulose BGCs (type I-III and acetylated cellulose) as well as a generic rule for cellulose BGCs which do not harbor any genes specific for a type.

**Making rules for single gene exoPS BGCs**

We also wanted to detect Sucrase-dependent BGCs. These pathways stand out, as they consist of a single membrane-bound sucrase enzyme which cleaves sucrose (or starch/maltodextrins[6]) and uses either glucose (alpha-glucansucrases) or fructose (fructansucrases) to assemble long polysaccharides[7,8]. Since epsProtocol was not designed for single gene systems, we manually created two Sucrase-dependent rules; one for alpha-glucansucrases and one for fructansucrases.

Alpha-glucansucrases belong to the glycohydrolase 70 (GH70) family, which belongs to the same clan (GH-H) as GH13 and GH77 in the CAZy classification system[9]. GH70 is the only member of the GH-H clan whose members are able to produce high-molecular weight alpha-D-glucans, using the D-glucopyranosyl donor obtained from cleaving sucrose, starch or maltodextrins[6,7]. Taking the amino acid sequences from a phylogenetic tree of representative GH70 sequences[6], we created a pHMM for alpha-glucansucrases in a manner identical to that in the epsProtocol (dereplication at 90% ANI followed by MAFFT alignment, trimming and hmmbuild). We then extracted all bacterial sequences with NCBI accessions from the GH-H clan as listed in the CAZy database and searched these with the newly created pHMM using hmmsearch. Sequence hits from the GH13 family all had bit scores at or below 125, while all GH77 sequence hits had bitscores at or below 22. The vast majority (95%) of GH70 sequences had a bit score above 200. The sequences below this score were all marked as either partial, hypothetical or both. We therefore set the bitscore threshold for this pHMM at 200 and added it as the alpha-glucansucrase rule in epsSMASH.

Fructansucrases belong to the glycohydrolase 68 (GH68) family, which are part of the same CAZy clan (GH-J) as GH32[9]. Fructansucrases catalyse the transfer of a fructosyl residue to a sucrose acceptor. The sucrose acceptor can vary *in vitro*, and includes water (sucrose hydrolysis), glucose (interchange reactions) and fructan (polymerisation)[8]. The majority of known fructansucrases are able to produce polysaccharides (levansucrase, inulosucrase, fructosyltransferase), however some only produce fructooligosaccharides (Beta-fructofuranosidase)[8,10]. We found a Pfam pHMM which was made to detect the GH68 family (*glyco_hydro_68*) and searched it against all bacterial sequences with NCBI accessions from the GH-J clan as listed in the CAZy database using hmmsearch. Sequence hits from the GH32 family all had bit scores at or below 31. In contrast, the vast majority (99%) of GH68 sequences had bit scores above 125. Most sequences assigned as GH68 below this score had bit scores below 70 and consisted of very small partial sequences (>100 aa). We therefore set the bitscore threshold for this pHMM at 125 and added it as the fructansucrase rule in epsSMASH.

**Creating relaxed and loose epsSMASH rules for the detection of novel putative exoPS BGCs**

The epsSMASH rules created using epsProtocol allowed us to detect known exoPS BGCs with well-described gene composition and synteny. Recently, an analysis of synthase-dependent exoPS BGCs with much looser detection requirements than our strict rules uncovered an alternate version of the pel operon in Gram-positive species[11]. To account for the possibility of such a discovery in any of the other synthase-dependent systems, we created less strict versions of all synthase-dependent exoPS BGC rules defined as "relaxed" rules (**Figure 1a**). These always required the synthase gene/complex and in every case but curdlan required another gene essential for exoPS secretion.

Wzx/Wzy- and ABC transporter-dependent systems differ from synthase-dependent systems by having a much more varied synteny and gene composition across different taxonomies. These systems include capsular polysaccharide and lipopolysaccharide pathways and are believed to be the most widespread methods of exoPS production[12]. Often, the genes of these systems are only conserved at a strain or species level[13]. Consequently, it was not possible to cover all described instances of these systems with our strict rules without inflating the ruleset to an unmanageable size. Instead, we exploit that core genes of these systems contain well-conserved domains, a strategy previously used to investigate polysaccharide export in diverse bacteria[14,15]. Based on the knowledge that all Gram-negative Wzx/Wzy- and ABC transporter-dependent systems require a polysaccharide co-polymerase (PCP) and an outer membrane polysaccharide export (OPX), we created a set of loose rules for the detection of putative Wzx/Wzy- and ABC transporter-dependent systems not caught by our strict rules. The most basic of these rules requires a PCP gene, an ABC-transporter or Wzx/Wzy gene, and at least two glycosyltransferases, to catch Gram-positive BGCs which do not contain an OPX gene. Stricter rules in the set require the OPX gene or genes specific to the Wzx/Wzy- or ABC transporter-dependent systems (Wzx, Wzy and ABC transporter proteins). The pHMMs used in the strict Wzx/Wzy- and ABC transporter-dependent rules as well as a collection of Pfam pHMMs were used to detect the different types of genes in these rules. The bit-scores of the pHMMs in these rules were set to 20. In the epsSMASH strictness hierarchy, these rules are in the "loose" category (**Figure 1a**).

**Supplementary Note 2: Validation of epsSMASH results**

Direct validation of epsSMASH BGC predictions would require a database of genomes with experimentally validated exoPS BGCs, which does not yet exist. Because of this, we made sure to perform a detailed manual analysis of each epsSMASH rule before implementing it, see Supplementary Note 1: "Testing pHMMs and creating epsSMASH rules". An additional validation step was performed by comparing the predictions of epsSMASH to previous attempts at predicting exoPS BGCs, namely the efforts of Bundalovic-Torma et al. (2020)[11] (**Supplementary Figure 1**) and Dueholm et al. (2023) (**Supplementary Figure 2**)[16] and by verifying epsSMASH predictions in model organisms (**Supplementary Table 2**)

Bundalovic-Torma et al. attempted to detect five synthase-dependent exoPS systems (cellulose, acetylated cellulose, alginate, pel and PNAG) across 1744 bacterial genomes retrieved from NCBI. They detected 4 acetylated cellulose, 64 alginate, 407 cellulose, 146 pel and 321 PNAG BGCs. Enabling the strict and relaxed rules and running epsSMASH on the 1744 genomes, we detect 88% (832/943) of these BGCs. The majority (~90%) of the additional hits detected by Bundalovic are 2-gene "gene clusters" or clusters with many duplicates of one gene if there are more than two. Some of them are not detected by the relaxed rules from epsSMASH due to the nature of the rule (e.g. our "pel-like" relaxed rule requires the synthase pelF and the export protein pelG, while some of the unique Bundalovic pel clusters consist of just the hydrolase pelA and pelF). Some of the missed BGCs would have been hit by a relaxed epsSMASH rule based on gene composition, so the reason they are not detected by epsSMASH is either because of a difference in our pHMMs or because our pHMM-specific bit scores threshold were more restrictive than their e-value threshold ($10^{-5}$). epsSMASH detects 7 cellulose and 2 pel BGCs which were not detected by the Bundalovic study. Three of these were detected by strict rules, and were classified as type 1 cellulose, generic cellulose and Gram-positive pel.

Dueholm et al. attempted to detect 16 different exoPS BGCs in a database of 1083 HQ MAGs obtained from activated sludge in Danish wastewater treatment plants. They detected 121 putative exoPS BGCs in as many genomes, classifying them as alginate, cellulose, hyaluronic acid, pel, PNAG, psl or xanthan BGCs. epsSMASH detects all the alginate and pel BGCs detected by Dueholm et al., as well as 7 pel BGCs not detected by them. A majority (31/44) of the cellulose BGCs detected by Dueholm et al. were not detected by epsSMASH. However, 30 of these predicted BGCs consisted of only a cellulose synthase (BcsA) homologue paired with either a transport protein (BcsC) or a glycohydrolase (bglX/BcsZ). According to Dueholm et al., these homologues exhibited percent identities near the 20% identity detection threshold. Based on their low sequence similarity and gene numbers, we conclude that these BGCs are likely not true cellulose systems. Notably, epsSMASH failed to detect one cellulose BGC and one PNAG-pga BGC, both of which contained more than two distinct genes. Manual inspection revealed that the genes in these BGCs were heavily fragmented, causing the bit scores of the epsSMASH pHMMs to fall below the defined thresholds for the epsSMASH rules. The hyaluronic acid, PNAG (eps), psl, and xanthan BGCs described by Dueholm et al. were not detected by epsSMASH. Dueholm et al. describe these BGC predictions as highly uncertain, since they did not follow the same synteny as described in literature. PNAG (eps), psl and xanthan are Wzx/Wzy-dependent systems, and the loose epsSMASH rules do detect at least one putative Wzx/Wzy-dependent BGC in each of the MAGs where Dueholm detects a PNAG (eps), psl or xanthan BGC.  We believe the comparisons above underscore epsSMASH's accuracy, as it detects likely genuine exoPS BGCs while avoiding highly uncertain predictions.

To estimate precision, we applied epsSMASH to the genomes of 14 well-characterized model organisms and manually verified the predictions. Among the 47 exoPS BGCs identified, only 3 were not described in literature, two of which were determined to be false positives, resulting in a positive predictive value of 96% (**Supplementary Table 2**). Although directly assessing epsSMASH's precision and sensitivity is challenging without a comprehensive database of experimentally confirmed exoPS BGCs, the results above suggest that the overall accuracy of the tool is high.

**Supplementary Note 3: Pel is a phylogenetically widespread pathway in activated sludge, the ocean and the human gut**

The pel pathway has been shown to be one of the most phylogenetically widespread exoPS BGCs in bacteria[17]. This is illustrated by the sequence similarity networks for MiDAS Global, GOMC and HumGut, all of which contain large BiG-SCAPE GCFs composed of pel BGCs found in species from 14 distinct phyla (**Figure 5**, **Supplementary Figure 7**). Despite this phylogenetic diversity, the synteny among the pel BGCs are highly conserved (**Supplementary Figure 7**). Pel operons from Pseudomonadota MAGs almost invariably display the canonical *pelABCDEFG* synteny, while operons from other Gram-negative phyla show alternative syntenies where *pelA*, *pelAB* or even *pelABC* are located after *pelDEFG* (**Supplementary Figure 7**). The pelDEFG proteins are believed to form an inner membrane complex which is responsible for pel polymerisation and transport into the periplasm[18], which likely explains why their synteny is conserved across phyla. Similarly, pelA and pelB form a complex which modifies and secretes periplasmic pel to the extracellular space[19], which could explain why their genes cluster together in most alternative pel syntenies.

Pel operons with the synteny *pelDEAFG* were detected by epsSMASH in Actinomycetota and Bacillota, the phyla where they were first described[20]. These operons lack the *pelB* and *pelC* genes which are localised in the periplasm and the outer membrane, and have so far been classified as Gram-positive pel systems. However, epsSMASH also detects several *pelDEAFG* operons in the Gram-negative phylum Chloroflexota, leading us to conclude that this operon is not exclusive to Gram-positive bacteria, but rather monoderm bacteria in general.

The Microflora Danica catalogue stood out as the only sequence similarity network in which pel exoPS BGCs were not present in the 20 largest BiG-SCAPE clusters. In fact, only 17 pel exoPS BGCs were detected in the entire MFD catalogue (**Supplementary Table 3**). Since the coverage of Microflora Danica core genera by the MFD catalogue exceeds 90% for all soil habitats[21], these results reveal that despite its ubiquity in many environments, pel is not a widespread system in Danish soils.

**Supplementary Note 4: Relaxed rules detect distant homologs to the pel operon**
In the MiDAS gene cluster similarity network, three BiG-SCAPE GCFs contained solely pel-like BGCs (**Figure 5**). The largest of these GCFs is made up of 48 pel-like BGCs found in genomes belonging to the Hyphomicrobiaceae family. The BGCs in this GCF are strongly conserved 6 gene operons beginning with a *pelA* gene and ending with *pelFG* (**Supplementary Figure 8**).

The second largest GCF contains 39 pel-like BGCs belonging to the phyla Pseudomonadota, Acidobacteriota, Chlamydiota and Verrucomicrobiota. The BGCs in this GCF are 6 gene operons beginning with *pelA* and ending with *pelDEFG*. A gene which epsSMASH fails to annotate is positioned between *pelA* and *pelD* and has a length similar to *pelB* from the *P. aeruginosa* pel operon.

The last BiG-SCAPE GCF contains 34 pel-like BGCs from the Verrucomicrobiales order. These are conserved 6-7 gene operons with the synteny *pelA?D?FG*, with question marks indicating genes not annotated by epsSMASH. Roughly half of the operons contain two unannotated genes between *pelA* and *pelD*.

Since all instances of pel-like operons had syntenies which resembled those of canonical, Gram-negative pel operons (a starting *pelA* and a concluding *pel*(*DE*)*FG*), and since Pfam HMMs detected tetratricopeptide-rich domains (found in *pelB* and *pelE*) and GAF domains (found in *pelD*) in many of the unannotated genes, we suspected that these were instances of distant *pelABCDEFG* homologs rather than novel pel operons such as the one found in Gram-positive pel producers[20].

To annotate the genes in these suspected distantly homologous operons, we performed structural homology searches of each protein in the three operons against both the PDB100[22] and the AlphaFoldDB[23] databases using Foldseek (v10.941cd33)[24]. Searches were carried out independently for each database using Foldseek's easy-search workflow, which integrates the creation of the database, structural similarity searching, and alignment conversion into a single pipeline. The --prostt5-model option was used to encode structural information using the 3Di alphabet. These searches confirmed that the unknown genes were homologs of pel proteins, namely pelB, pelD and pelE (**Supplementary Figure 8, Supplementary Table 4**). Curiously, none of the pel-like operons contained homologs of pelC, an outer membrane-associated protein which is thought to guide pel towards the pelB secretion channel[25].

Having confirmed the identities of the unannotated pel genes, we created new Hidden Markov Models for *pelB*, *pelD* and *pelE*. This allowed us to annotate all genes in the three pel-like operons, confirming that they are indeed distantly homologous versions of the canonical *pelABCDEFG* operon. The above process highlights the utility of relaxed epsSMASH rules for identifying distant homologs of known synthase systems by searching for conserved core genes within these systems.
